# A Scoring System to Evaluate the Impact of SNPs in a Path Related Context to Study Behçet’s Disease Aetiology in Japanese Population

**DOI:** 10.1101/316562

**Authors:** Ozan Ozisik, Akira Meguro, Nobuhisa Mizuki, Banu Diri, Osman Ugur Sezerman

## Abstract

**Motivation:** Genome-wide association study (GWAS) is a powerful method that can provide a list of single nucleotide polymorphisms (SNPs) that are significantly related to the pathogenesis of a disease. Even though in Mendelian diseases strong associations can be found for certain SNPs, in most of the complex diseases only modest associations can be identified from the GWAS. Therefore, the main challenge in such studies is to understand how multiple SNPs that have modest association with the phenotype interact and contribute to its aetiology. This can only be done via pathway based analysis of modestly associated SNPs and the genes that are affected by these changes.

**Results:** In this study, we propose DAPath, a Disease Associated Path analyzer tool for discovering signaling paths and the pathways that contain these paths which are subjected to cumulative impact of modestly associated variants. We applied our proposed method on Behçet’s disease (BD) GWAS dataset from Japanese population. Antigen Processing and Presentation pathway is ranked first with 16 highly affected paths. Th17 cell differentiation, Natural killer cell mediated cytotoxicity, Jak-STAT signaling, and Circadian rhythm pathways are also found to be containing highly affected paths.

**Availability:** The proposed method is available as a Cytoscape plug-in through https://github.com/ozanozisik/DAPath

## 1. Introduction

Behçet’s disease (BD) is a chronic immune-mediated disease which affects multiple parts of the body. Common manifestations are ocular lesions, oral aphthosis and genital aphthosis. A genetic tendency to uncontrolled inflammatory reactions induced by various environmental triggers is considered to play a critical role in the development of BD. Association between BD and HLA-B51 has been known for a long time and confirmed in different studies (de Menthon et al. 2009). Two landmark genome-wide association studies (GWAS) on Japanese and Turkish populations (Mizuki et al. 2010; Remmers et al. 2010) have identified HLA-A, IL10, and IL23R–IL12RB2 to be novel BD susceptibility loci, in addition to confirming HLA-B51. In (Xavier et al. 2013) 508 genes have been found to be differentially expressed which have been used for further pathway analysis and association study. In (Bakir-Gungor et al. 2015), pathway analysis has been performed on GWAS datasets from Turkish and Japanese populations and common pathways found to be important despite different SNP affected genes.

Although pathway analysis gives a level of understanding for affected processes in a disease, it is necessary to identify affected set of interactions within a pathway to understand disease mechanisms. This will also lead to the way to individualized treatment decisions.

In the last decade, there have been many integrative studies that overlay interaction networks with molecular profiles, e.g. differential expression data or GWAS data to identify “active modules”. Active module identification is categorized into three classes by Mitra et al. (2013): i) active subnetwork search methods (also known as significant area search), ii) biclustering methods, and iii) network propagation methods. In active subnetwork search methods (Ideker et al. 2002; Chuang et al. 2007; Dittrich et al. 2008; Ulitsky and Shamir 2009; Ma et al. 2011; Doungpan et al. 2016; Ozisik et al. 2017), it is aimed to find connected subgraphs of genes that contribute to the disease condition collectively in a biological interaction network. In biclustering (Cheng and Church 2000; Hanisch et al. 2002; Reiss et al. 2006), it is aimed to cluster network interactions and the conditions under which these interactions are active. In active subnetwork search and biclustering methods, a list of genes related to the condition is obtained but these methods do not clarify the impact on signal flow cascades. In network propagation methods (Draghici et al. 2007; Tarca et al. 2009; Vaske et al. 2010; Dutta et al. 2012; Haynes et al. 2013; Sebastian-Leon et al. 2014; Li et al. 2015; Bokanizad et al. 2016), molecular profiles are used with directed gene networks to determine their effects on the signal flow. The advantage of network propagation methods over other methods is considering the interactions among genes to understand how an alteration affects signaling taking into account the position of the signature gene in the network.

Draghici et al. presented the seminal study that uses network propagation (Draghici et al. 2007). This method is improved by the same group in (Tarca et al. 2009) and Signaling Pathway Impact Analysis (SPIA) method is proposed. In SPIA, two types of evidence are combined to determine whether a signaling pathway was impacted by observed changes: i) overrepresentation analysis of the number of differentially expressed genes, ii) the abnormal perturbation of the pathway caused by the position of the differentially expressed genes and connections in the pathway. In (Vaske et al. 2010), a factor graph is created for each gene; entities in the graph represent the copy number of the genome, mRNA expression, protein level, and protein activity, allowing the incorporation of many types of omic data as evidence. The method predicts the degree to which a pathway’s activities are altered using probabilistic inference. In PathNet proposed in (Dutta et al. 2012), KEGG pathways are combined to create an interaction network. Each gene’s score based on expression data is combined with its neighborhood score using the Fisher’s method. The neighborhood score is calculated by adding the logarithms of its neighbors’ scores and then calculating the obtained score’s significance using a bootstrap approach. Related pathways are obtained using hypergeometric test. In (Haynes et al. 2013), path with maximum running sum score is found for each pathway. Rotation test is used to infer the significance. The maximum scoring path is used to test the null hypothesis about the expression of the entire pathway. Sebastian-Leon et al. (2014) proposed PATHiWAYS that handles pathways as collections of stimulus-response circuits containing alternative paths that lead an input node to an output node. Each circuit’s activation probability is calculated and the number of significantly activated circuits is compared between two conditions. Gene expression measurements are transformed to node probabilities. A circuit’s activation probability is calculated by multiplying probabilities of nodes on the same path and calculating the probability of activation of at least one path. Li et al. (2015) proposed Sub-SPIA method that applies SPIA to subpathways. These subpathways are acquired by starting with significant nodes in the pathway, connecting the ones that have at most n non-significant genes in-between, applying Kruskal algorithm to find minimum spanning tree and removing unnecessary non-significant genes. A pathway is altered if one of its paths is significantly altered. In (Bokanizad et al. 2016), SPIA is improved by considering inter-pathway interactions while calculating perturbations.

In this study, we propose DAPath, a disease associated path analyzer tool for discovering signaling paths under pathways that are subject to cumulative impact of modestly associated variants. With pathway analysis in signaling path level, we can find the most affected interaction cascades and potential drug targets that may counteract these alterations. We applied our proposed method on Behçet’s disease (BD) GWAS dataset from Japanese population. We compared our results with the results of SPIA and Sub-SPIA.

## 2. Materials and Methods

### 2.1. Datasets

We applied our method to Behçet’s disease (BD) GWAS dataset from Japanese population. Mizuki et al. (2010) conducted GWAS on 612 Japanese individuals with BD and 740 healthy controls. The list of significant genes was obtained using PANOGA by Bakir-Gungor et al. (2015).

We used pathway data from Kyoto Encyclopedia of Genes and Genomes (KEGG) repository (Kanehisa and Goto 2000; Kanehisa et al. 2016). The download date of the pathways is April 23rd, 2017. Some small corrections have been made on the provided KEGG XML files which contradict with the provided graphics, e.g. a missing connection between nodes is added, a group of nodes that are not tagged accordingly are tagged as a group in the KEGG XML file. We discarded metabolic pathways since our focus was not on metabolic diseases and our dataset did not contain metabolite information.

### 2.2. Proposed Path Impact Analysis Method

In this study we propose a network propagation method to analyze the impact of genes and their significance values on cascades of interactions (signaling paths) in KEGG pathways. A path in a pathway is composed of an input node, an output node, and the intermediate nodes that constitute the path in-between. If there are alternatives in-between, each is accepted as a distinct path. In the proposed method, significance values of nodes are preprocessed considering their crosstalk and whether they are part of a complex; then impact score for each path is calculated and affected pathways are determined from the significance scores of altered signaling paths that the pathways contain.

#### 2.2.1. Crosstalk Treatment

Donato et al. (2013) showed that crosstalk can cause a biologically non-significant pathway become statistically significant and vice versa in enrichment applications. In order to prevent common genes with high scores from leading to many high scoring irrelevant paths, we decreased the significance of genes that belong to multiple pathways. In crosstalk treatment it is needed to determine crosstalk tolerance and crosstalk factor range. Crosstalk tolerance is the threshold for the number of pathways a gene can be member of without being subject to treatment. Crosstalk factor is the multiplier used to increase p-value and crosstalk factor range is the range that crosstalk factor will change in proportion to the number of pathways the gene belongs to.

#### 2.2.2. Significance of Nodes

In genome-wide association studies, p-value threshold of 5 × 10^−8^ has become a standard (Fadista et al. 2016) but this threshold causes the elimination of the genes with modest association to the condition that might have aggregate effects on the pathways they are involved. In this study we used SNPs with nominal evidence of association (p-value<0.05) following the approaches in (Baranzini et al. 2009) and (Bakir-Gungor and Sezerman 2011).

For each gene, if the gene is in the dataset, we assigned its p-value as its significance, otherwise we assigned a base p-value, 0.05, which is the threshold to accept SNPs in this study (Eq. (1)).

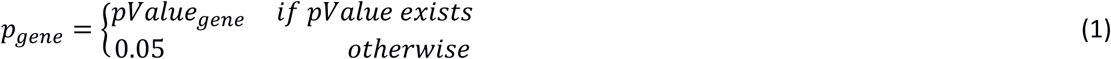

In KEGG pathways, there are nodes that are composed of multiple alternative genes. There are also gene groups which correspond to complexes of gene products, mostly protein complexes. The minimum p-value of multiple genes assigned to a node is taken as the p-value of that node (Eq. (2)).

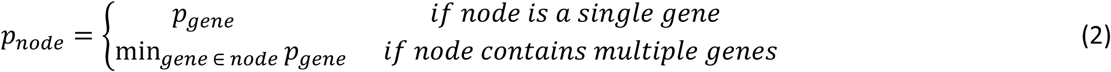

#### 2.2.3. Impact Score Calculation

A path’s impact p-value is calculated by multiplying p-values of nodes on it (Eq. (3)). In some pathways (e.g. Jak-STAT signaling pathway) one gene is represented in two consecutive nodes and connected by state change. In this condition, the gene’s p-value is used only once.

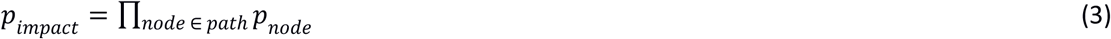

Since we are multiplying p-values in Eq. (3), longer paths will have stronger significance values. We normalized the scores to compensate for these effects. We calculated a base p-value for each path by assigning the base node p-value, 0.05, to all nodes (Eq. (4)), and calculated each path’s impact score using Eq. (5). This score is close to 1 when the base p-value and the path p-value are close to each other, and higher when the path contains significant genes.

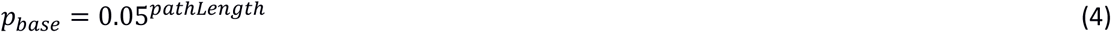

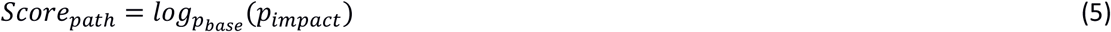

### 2.3. Comparative study

The studies in this domain, SPIA, Sub-SPIA and PATHiWAYS, are designed for expression changes but we aimed to see if they are applicable to GWAS data and compare with our method. PATHiWAYS method takes Affymetrix Human Genome microarray data as input. Our study is on GWAS data. PATHiWAYS cannot be applied to all pathways, some pathways are densely connected and there are thousands of alternative paths between an input and an output node, calculating the probability of the union of these paths is not feasible. PATHiWAYS discard circuits containing loops. Hence we could not use PATHiWAYS for comparison. We compared our method with SPIA and Sub-SPIA. These methods use log fold changes as input. Our data consists of GWAS p-values. Log base 10 of p-values are used to get values that are compatible with SPIA and Sub-SPIA.

## 3. Results and Discussion

We applied our method on Behçet’s disease GWAS dataset.

The parameters of the method are related to crosstalk treatment. We performed trials using different values for crosstalk tolerance and crosstalk factor. We tried {1, 2, 3, 4, 5} as crosstalk tolerance. For crosstalk factor range we tried [10, 100] with an increment of 5, [10, 100] with an increment of 10, [10, 1000] with an increment of 10. We saw that the method is not oversensitive to these parameters and rank of the high scoring pathways change a few steps up or down. Finally we set crosstalk tolerance to 3 and crosstalk factor range to [10, 100] with an increment of 5.

Affected pathways are given in Table 1. Pathways are sorted by the score of the highest scoring path they contain. Pathways in the literature are written in boldface.

**Table 1.** Pathways that contain the highest scoring disrupted paths

| No | Pathway | Score | Number of Paths |
| --- | --- | --- | --- |
| 1 | Antigen processing and presentation | 5.48 | 16 |
| 2 | Type II diabetes mellitus | 3.46 | 2 |
| 3 | Maturity onset diabetes of the young | 3.08 | 4 |
| 4 | Th17 cell differentiation | 2.96 | 11 |
| 5 | Natural killer cell mediated cytotoxicity | 2.93 | 60 |
| 6 | Focal adhesion | 2.72 | 545 |
| 7 | Tuberculosis | 2.67 | 9 |
| 8 | Jak-STAT signaling pathway | 2.38 | 20 |
| 9 | MAPK signaling pathway | 2.29 | 240 |
| 10 | Circadian rhythm | 2.25 | 5 |
| 11 | Osteoclast differentiation | 2.23 | 3 |
| 12 | Insulin resistance | 2.20 | 2 |
| 13 | Aldosterone synthesis and secretion | 2.20 | 1 |
| 14 | Axon guidance | 2.17 | 1 |
| 15 | PI3K-Akt signaling pathway | 2.16 | 2 |
| 16 | Phosphatidylinositol signaling system | 2.15 | 13 |
| 17 | Dopaminergic synapse | 2.06 | 2 |
| 18 | Insulin signaling pathway | 1.98 | 1 |
| 19 | p53 signaling pathway | 1.98 | 1 |
| 20 | FoxO signaling pathway | 1.97 | 1 |

Our method does not apply a threshold of significance or score to the paths, it provides an interface for the researchers to be able to observe each affected path. Even in enrichment studies, determining a threshold for the pathways to present is difficult. In the results of SPIA 117 pathways have p-value less than 0.05. We chose to present the first 20 pathways and their affected paths that have score above the 21st pathway. Figures of the affected paths are available as supplementary file.

Some of the important paths found to be disrupted by our method are given below. While mentioning genes, symbols given on KEGG visuals are given first, gene names are given in parenthesis if they are different. Genes’ p-values (after crosstalk treatment) are indicated using asterisk symbol, single asterisk (*) for p-values in [0.01, 0.05] range, double asterisk (**) for p-values in (5 × 10^−8^, 0.01) range, and triple asterisk (***) for more significant genes.

Antigen processing and presentation contains the most affected paths. Affected paths on MHCI pathway start from CIITA (*), IFNγ (IFNG*), TNFα (TNF) and lead to KIR (KIR2DS1***, KLRC1*, KLRC2**, KLRD1*, KLRC4**) and CD8 (CD8B*) through MHCI (HLA-A**, HLA-B***, HLA-C***, HLA-E*, HLA-F***, HLA-G***). These paths affect natural killer cells and cytotoxic T cells. Affected paths on MHCII pathway start from CIITA (*), CTSB (*), CTSS (*) and lead to CD4 (*) though MHCII (HLA-DMB*, HLA-DOA*, HLA-DOB*, HLA-DQA1*, HLA-DQA2**, HLA-DQB1***, HLA-DRA**). In (Oguz et al. 2016), it is concluded that negative regulation of antigen processing and presentation proteins (CTSS, ERAP1) appear to be instrumental in Behçet’s Disease immunopathogenesis.

Natural killer cell mediated cytotoxicity pathway is also highly affected, the disrupted paths start from HLA-C (***), continue with KIR2DS (KIR2DS1***) and lead to cytokine release by TNFα (TNF) and GM-CSF (CSF2*), and cytotoxic granule release by PKC (PRKCA*, PRKCB*).

Antigen processing and presentation pathway and Natural killer cell mediated cytotoxicity pathway are among the top affected pathways in our study and in both of them TNFα takes part. TNFα is a therapeutic target. Anti-TNFα therapy has been declared as effective on BD in many studies (Arida 2011; Calvo-Rio et al. 2014; Vallet et al. 2015).

In Th17 cell differentiation pathway, IL-23 (IL23A*) – IL23R (**) – JAK2 (*), TYK2 (*) – STAT3 path that again regulates IL23R (**) is the most affected path. The same path also regulates IL17A, IL17F, IL21 (*), IL22, HIF1A, RORA (*), RORγt (RORC). This path affects protection against extracellular pathogens and autoimmunity. The role of Th17 Cells in the pathogenesis of BD has been reviewed in (Nanke et al. 2017). Another path that regulates IL23R, TGFβ (TGFB1) – TGFβ-R (TGFBR2*) – SMADs (SMAD2*, SMAD3*, SMAD4) – IL23R path is also affected.

In Maturity onset diabetes of the young pathway the affected paths start with HNF6 (ONECUT1*), Hex (HHEX*) or Hlxb9 (MNX1) and continue with PDX1 and NR5A2 (***). In (Venteclef et al. 2010) anti-inflammatory role of NR5A2 (referred as LRH-1) has been stated. In (Venteclef et al. 2006) it is shown that NR5A2 is a key player in the control of the hepatic acute-phase response. In (Coste et al. 2007) it is shown that NR5A2 has an important role in controlling of intestinal inflammation and the pathogenesis of inflammatory bowel disease. In KEGG the outcome of the affected paths are not clear yet.

In Focal adhesion pathway, paths leading to CycD (CCND1, CCND2, CCND3), BAD and BCL2 are affected. These paths affect cell proliferation and cell survival. This pathway is found to be important in both Japanese and Turkish populations by Bakir-Gungor et al. (2015).

In Jak-STAT signaling pathway, paths through Cytokines (CSF2*, IFNB1*, IFNG*, IFNW1*, IL7**, IL10**, IL12A*, IL15*, IL22*, IL23A*, IFNK*, IL21*, TSLP**), Receptors (CSF2RB*, CSF3R*, IL23R**, IFNLR1**, IFNAR1*, IFNAR2*, IFNGR1*, IFNGR2*, IL2RA*, IL2RB*, IL4R*, IL5RA**, IL6R*, IL10RA*, IL12RB2*, LEPR*, LIFR*, IL20RB**, PRLR*, IL22RA1**, OSMR**), JAK (JAK2*, TYK2*), STAT (STAT1*, STAT4*, STAT5A*) and target genes are affected. This pathway’s activation in BD has been demonstrated by Tulunay et al. (2015).

In MAPK signaling pathway, paths starting with CACN (CACNA1E***) and leading to phosphorylation of Tau (MAPT), STMN1, cPLA2 and c-Myc are highly disrupted. MAPT encodes the microtubule-associated protein tau which stabilizes microtubules. STMN1 is involved in the regulation of the microtubule filament system by destabilizing microtubules. Colchicine, a microtubule inhibitor is often used in treatment of BD (Shelef et al. 2013). In (Saga et al. 1987), it is reported that the distribution of microtubules in the neutrophils of BD patients and controls were different which suggested that microtubules played some significant role in the pathophysiology of BD.

In Circadian rhythm pathway, paths leading through Per, Cry (CRY2**), CSNK1, Dec (BHLHE40**), Bmall (ARNTL*) and Clock (NPAS2*) are highly disrupted. This pathway controls clock output and rhythmic biological processes. In (Hand et al. 2016), disruption of the circadian clock is stated as an aggravating factor associated with a range of human inflammatory diseases. Yu et al. (2013) showed the role of circadian clock on Th17 cell differentiation.

In Osteoclast differentiation pathway, FHL2 (**) – TRAF6 (*) – TAK1 (MAP3K7**) – TAB1/2 – MEK1 (MAP2K1*) and FHL2 (**) – TRAF6 (*) – TAK1 (MAP3K7**) – MKK6 (MAP2K6*) – p38 (MAPK13*, MAPK14*) – AP1 (FOSL2*) – CTR (CALCR**) paths are affected. In (Ciofani et al. 2012) it is found that FOSL2 limits plasticity of T helper cell differentiation and plays a key role in a mouse model of autoimmune disease. In (DuPage and Bluestone 2016) it is stated that T cell plasticity is an important factor in immunological diseases. In (Komatsu et al. 2014) contribution of plasticity of IL-17^+^FOXP3^+^ Treg cells towards T_H_17 cells to the pathogenesis of rheumatoid arthritis has been shown. In (Cafforio et al. 2009) it is suggested that CALCR may contribute to the modulation of cytoplasmic calcium(2+) levels needed to regulate T and B cell activation and perhaps other immune functions. These paths are also modulated by TNFα which is a therapeutic target as stated before.

In Insulin resistance pathway, PTPs (PTPRF**) – IRS-1** – PI3K (PIK3R1*) – PKCζ (PRKCZ) path which leads to inhibition of GLUT4 translocation and glucose uptake is disrupted. Insulin resistance has been associated with BD in (Erdem et al. 2006) and (Kim et al. 2010).

In Axon guidance, TRPC (TRPC3**, TRPC4**, TRPC6*) – CaN (PPP3CA*)– NFAT (NFATC2*) path leading to axon outgrowth is disrupted. In (Birol2004), peripheral nerve involvement in BD has been mentioned and it has been reported that peripheral nerve dysfunction of BD is an axonal type of distal polyneuropathy that predominantly involves the lower extremities.

We compared our results with the results of Signaling Pathway Impact Analysis (SPIA) method proposed by Tarca et al. (2009) and Sub-SPIA method proposed by Li et al. (2015). In Table 2 and Table 3, most significant 20 pathways found by SPIA and Sub-SPIA are given respectively.

**Table 2.** Results of SPIA. First 20 pathways sorted by P_G_ are given.

| No | Pathway | $P_{NDE}$ | $P_{PERT}$ | $P_G$ |
| --- | --- | --- | --- | --- |
| 1 | Pathways in cancer | 7.07E-26 | 7.53E-01 | 3.15E-24 |
| 2 | Neuroactive ligand-receptor interaction | 5.46E-23 | 6.57E-01 | 1.89E-21 |
| 3 | MAPK signaling pathway | 1.13E-17 | 4.93E-01 | 2.28E-16 |
| 4 | Calcium signaling pathway | 4.57E-17 | 1.23E-01 | 2.29E-16 |
| 5 | Axon guidance | 1.80E-16 | 8.20E-02 | 5.88E-16 |
| 6 | Focal adhesion | 1.44E-16 | 3.05E-01 | 1.70E-15 |
| 7 | Vascular smooth muscle contraction | 8.04E-16 | 1.22E-01 | 3.71E-15 |
| 8 | Toxoplasmosis | 6.23E-16 | 7.99E-01 | 1.80E-14 |
| 9 | Serotonergic synapse | 2.44E-14 | 5.90E-02 | 5.06E-14 |
| 10 | Cytokine-cytokine receptor interaction | 1.08E-13 | 9.60E-02 | 3.43E-13 |
| 11 | Glutamatergic synapse | 4.49E-14 | 4.12E-01 | 6.04E-13 |
| 12 | Retrograde endocannabinoid signaling | 7.64E-14 | 3.69E-01 | 9.08E-13 |
| 13 | Jak-STAT signaling pathway | 3.62E-12 | 3.78E-01 | 3.87E-11 |
| 14 | Cholinergic synapse | 3.82E-10 | 5.00E-03 | 5.34E-11 |
| 15 | ECM-receptor interaction | 9.49E-12 | 6.39E-01 | 1.63E-10 |
| 16 | Antigen processing and presentation | 3.01E-06 | 5.00E-06 | 3.90E-10 |
| 17 | HTLV-I infection | 1.62E-11 | 9.45E-01 | 3.96E-10 |
| 18 | Dopaminergic synapse | 4.72E-11 | 7.36E-01 | 8.71E-10 |
| 19 | Gap junction | 4.20E-10 | 9.00E-02 | 9.44E-10 |
| 20 | TGF-beta signaling pathway | 3.29E-10 | 3.19E-01 | 2.52E-09 |

**Table 3.** Results of Sub-SPIA. First 20 pathways sorted by P_G_ are given.

| No | Pathway | pNDE | pPert | pG |
| --- | --- | --- | --- | --- |
| 1 | MAPK signaling pathway | 2.69E-26 | 3.42E-01 | 5.61E-25 |
| 2 | PI3K-Akt signaling pathway | 2.16E-24 | 5.95E-01 | 7.18E-23 |
| 3 | Focal adhesion | 1.60E-19 | 4.78E-01 | 3.44E-18 |
| 4 | Ras signaling pathway | 4.17E-18 | 1.18E-01 | 2.12E-17 |
| 5 | Rap1 signaling pathway | 4.86E-18 | 8.02E-01 | 1.60E-16 |
| 6 | Pathways in cancer | 3.70E-17 | 8.26E-01 | 1.19E-15 |
| 7 | cAMP signaling pathway | 1.12E-15 | 2.49E-01 | 1.02E-14 |
| 8 | Neurotrophin signaling pathway | 1.21E-13 | 1.11E-01 | 4.44E-13 |
| 9 | Osteoclast differentiation | 3.55E-14 | 6.23E-01 | 7.18E-13 |
| 10 | Tuberculosis | 6.79E-14 | 6.77E-01 | 1.46E-12 |
| 11 | p53 signaling pathway | 1.04E-13 | 4.99E-01 | 1.64E-12 |
| 12 | Oxytocin signaling pathway | 3.76E-13 | 3.50E-01 | 4.03E-12 |
| 13 | Gap junction | 7.53E-13 | 2.91E-01 | 6.61E-12 |
| 14 | cGMP-PKG signaling pathway | 5.98E-13 | 5.29E-01 | 9.43E-12 |
| 15 | Dopaminergic synapse | 4.58E-13 | 7.00E-01 | 9.54E-12 |
| 16 | Jak-STAT signaling pathway | 1.31E-11 | 9.30E-02 | 3.46E-11 |
| 17 | Wnt signaling pathway | 1.34E-11 | 1.41E-01 | 5.28E-11 |
| 18 | Calcium signaling pathway | 2.52E-12 | 9.81E-01 | 6.85E-11 |
| 19 | TGF-beta signaling pathway | 2.50E-11 | 5.44E-01 | 3.54E-10 |
| 20 | Autophagy - animal | 7.67E-11 | 4.83E-01 | 9.26E-10 |

In SPIA, a very important pathway, Antigen processing and presentation, is 16th according to P_G_, this is the result of relatively big P_NDE_. This pathway is the most significant pathway according to P_PERT_. In SPIA, Natural killer cell mediated cytotoxicity is not in top 20 according to P_G_, but it is 2nd according to P_PERT_. We see that SPIA results are dominated by overrepresentation analysis results. Jak-STAT signaling pathway and Focal adhesion are the examples of the opposite case where the pathways are significant according to P_NDE_, which lets them in top 20 according to P_G_, but they are not significant according to P_PERT_. Th17 cell differentiation, Osteoclast differentiation and Circadian rhythm pathways are not among the first 20 pathways in any case. In Sub-SPIA, Antigen processing and presentation, Natural killer cell mediated cytotoxicity, Th17 cell differentiation, Circadian rhythm, Insulin resistance, and Axon guidance pathways which are found to be affected by DAPath are not in the top 20 pathways. One missing pathway in DAPath is TGF-beta signaling pathway, which is known to be related to Behçet’s disease (Bakir-Gungor et al. 2015). This pathway is among the top 20 pathways in both SPIA and Sub-SPIA.

## 4. Conclusion

In this study, we proposed DAPath, a method to analyze the cumulative effect of variants on signaling paths. This approach enables us to infer which outcomes of the pathway are affected, understand disease mechanisms, and identify possible drug targets.

We applied our method on Behçet’s disease GWAS data and our method extracted important paths in pathways related to inflammation and immunity, e.g. Antigen processing and presentation, Th17 cell differentiation, Natural killer cell mediated cytotoxicity, Focal adhesion, MAPK signaling pathway, Jak-STAT signaling pathway. The results also show the cumulative importance of the modestly associated genes rather than single variations in cascades of interactions. Circadian rhythm pathway, Osteoclast differentiation pathway, Insulin resistance pathway and Axon guidance pathway are good examples of the aggregate effect of modestly associated genes. None of the genes on the disrupted paths of these pathways have a p-value less than 5 × 10^−8^, which is the threshold used in GWAS. These pathways would be missed in a standard GWAS analysis.

